# Functional assessments of short-term spatial memory in the Dog Aging Project identify strong associations with age that are not moderated by body mass

**DOI:** 10.1101/2025.06.30.662397

**Authors:** Stephanie H. Hargrave, Amber J. Keyser, Emma Kristal, Gene E. Alexander, Theadora A. Block, Emily E. Bray, Laura E. L. C. Douglas, Brenda S. Kennedy, Daniel E. L. Promislow, David A. Raichlen, Dog Aging Project Consortium, Evan L. MacLean

## Abstract

Companion dogs have emerged as a valuable model in the study of cognitive aging, but assessments of cognitive function in large, diverse, and geographically distributed samples of dogs are challenging to obtain. We developed two novel functional assessments of short-term spatial memory that were administered by community science participants in a sample of 6,753 dogs through the Dog Aging Project. We compared data generated by community scientists to those gathered by research professionals, estimated relationships between age and task performance, and tested the hypothesis that associations between age and cognitive performance vary by dog body mass, as a proxy for expected lifespan. Community scientists generated similar data to research professionals and both cognitive tasks were sensitive to age-related deficits, beginning in midlife. Relationships between age and cognitive function were highly similar across small and large dogs and, for both tasks, comparison of models with and without an interaction between age and body mass yielded decisive evidence for the model without the interaction. Large dogs exhibit accelerated aging across many traits, and so the lack of evidence for accelerated cognitive aging raises the possibility that their large size confers a neuroprotective advantage. We consider possible mechanisms underlying this effect and address how experimental studies of dog cognition using community science methods can support future research on mechanisms of brain and cognitive aging.

## Introduction

Companion dogs have emerged as an important model in aging research; they share a wealth of features with humans while also having comparatively shorter lifespans, which enables efficient longitudinal research (Gilmore & Greer, 2015; Ruple et al., 2021). Unlike laboratory animals, companion dogs share their environment with humans, have access to a sophisticated healthcare system, and are genetically diverse (Kaeberlein et al., 2016). Dogs also develop many of the same age-related diseases as humans, with dogs and humans often sharing similar genetic and environmental risk factors (Pallotti et al., 2022).

Studies of cognitive aging in dogs have identified parallels with human cognitive decline, including age-related deficits in learning and memory that often begin in midlife (Chapagain et al., 2018; Head, 2013). Some senior dogs also spontaneously progress to a profoundly impaired state, termed canine cognitive dysfunction, with some behavioral and neuropathological similarities to Alzheimer’s disease (González-Martínez et al., 2011; Head, 2011; Osburn et al., 2024; Prpar Mihevc & Majdič, 2019; Urfer et al., 2021). Although the potential value of dogs as a model for human cognitive decline, dementia, and specific neurodegenerative diseases such as Alzheimer’s disease has become widely recognized, the full benefits of a companion dog model have yet to be realized due to the challenges of large-scale research with this population.

To date, studies of cognitive aging in dogs have been conducted primarily in two distinct research traditions, each with strengths and weaknesses. The first approach is a traditional laboratory model in which a relatively small and intensively studied population is maintained and studied under laboratory conditions. Research following this model was central to initial discoveries regarding age-related cognitive dysfunction in dogs and potential neurobiological similarities to Alzheimer’s disease (Adams et al., 2000; Head, 2013). Traditional laboratory studies have benefited from rigorous experimental approaches to the study of cognition, employing objective tests of learning and memory as primary cognitive endpoints. However, such studies have typically been conducted with small samples of a single breed (beagles) studied under laboratory conditions and thus have not capitalized on several important features of companion dogs, including genetic diversity, variable and shared environments with humans, and the potential for large-scale research in the natural environment. A second approach has prioritized the study of large, genetically and phenotypically diverse samples of dogs living in human environments. Research in this tradition has been critical for the epidemiology of dementia in dogs, including estimates of disease prevalence and our emerging understanding of risk factors shared with humans (Azkona et al., 2009; Salvin et al., 2010). However, when working with large and geographically distributed samples, it has not been possible to measure cognitive function using experimental approaches. Rather, most studies in this tradition have used owner-report surveys that ask about behavioral signs of dementia. Although several such scales have been validated for detection of canine cognitive dysfunction (Madari et al., 2015; Salvin et al., 2011), these assessments rely on dog owners to notice and relay behavioral changes associated with dementia, and the relative subjectivity of doing so has numerous potential pitfalls.

More recently, some scientists have explored a third approach in which pet owners administer simple cognitive assays in the home environment (Hoel et al., 2021; Pelgrim et al., 2024; Stewart et al., 2015; Watowich et al., 2020). To facilitate this method, scientists have designed simplified cognitive tests that can be completed using commonly available household materials. A key aim of this approach is to enable objective functional assessments of cognition while also capitalizing on advantages associated with the large-scale study of privately owned companion dogs. Initial research using this approach has demonstrated that community scientists can produce data similar to those obtained in laboratories (Stewart et al., 2015), performance in home-based tests is repeatable and associated with clinical measures of dementia (Hoel et al., 2021), and age-related variation can be captured in large samples (Watowich et al., 2020). Though initial studies have been promising, the experimental study of dog cognition using community science is a nascent area of research, and the development of assays tailored for studies of aging are critical for future success.

In the current work, we developed two novel assessments to measure aspects of spatial memory associated with cognitive decline and deployed these assessments in the Dog Aging Project, a nationwide study of aging in U.S. companion dogs (Creevy et al., 2022). Using data from these measures we modeled relationships between age and cognitive performance and evaluated whether body mass, a strong predictor of lifespan in dogs, moderates this relationship. The shorter lifespans of large dogs have been attributed to faster rates of aging (Kraus et al., 2013), evidence for which has been observed across a wide range of biological processes (McCoy et al., 2024). However, whether the timing of cognitive decline relates to body mass in dogs remains poorly understood. Whereas some studies have identified different size-associated trajectories (Turcsán & Kubinyi, 2024), others have not (Watowich et al., 2020). Lack of a moderating role for body size in cognitive aging would suggest that larger dogs maintain cognitive health for a greater fraction of their expected lifespan, potentially providing insights into mechanisms that maintain brain and cognitive health, despite declines in other body systems. In contrast, evidence that cognitive decline occurs earlier and/or more rapidly in larger dogs would yoke these phenomena more closely to other patterns of age-related decline, potentially pointing to more systemically integrated mechanisms underlying cognitive health.

## Methods

### Subjects and Recruitment

The Dog Aging Project is a longitudinal open-science study of aging in a nationwide sample of companion dogs in the United States. Participating dogs are nominated by their owners, who provide annual information about their dog’s health, environment, and lifestyle. The overall structure of the study and its sampled cohorts are described in Creevy et al. (2022). Data in this manuscript reflect participating dogs’ first exposure to two cognitive assessments, 1-2-3 Treat and Treat Hide & Seek) associated with 2023 and 2024 project data releases (Dog Aging Project, 2024, 2025). All members in the DAP Pack were invited by email to complete 1-2-3 Treat in February of each year and Treat Hide & Seek in August of each year. This email contained a link to their personal research portal where they had the opportunity to complete the activity or opt-out of the activity and explain why it was not a good fit for them and/or their dog. The activity remained open for 40 days from the date of invite. Owners received three reminder emails at day 10, day 20, and day 30 if they had not already completed the activity.

Demographic information for participating dogs is shown in Table 1. To restrict analyses to adult animals, data from dogs < 1 year of age were excluded. We also excluded data from dogs reported to be > 18 years of age. The size distribution of dogs that participated in the cognitive tasks was similar to that for non-participant dogs, but dogs that participated in the cognitive activities tended to be moderately younger than non-participants (Supplemental Materials). Additionally, owners of participating dogs tended to be older than owners of non-participant dogs (Supplemental Materials). Study-related procedures involving these privately-owned dogs were approved by the Texas A&M University IACUC, under AUP 2021-0316 CAM. To compare data collected by community scientists with those collected in conventional research settings, we recruited an independent sample of 30 companion dogs (14 female, 16 male, age (mean ± SD = 8.1 ± 4.2 years), weight (mean ± SD = 22 ± 10 kg)) that were tested at the Arizona Canine Cognition Center, or nearby satellite sites, by research professionals using the same protocols (University of Arizona IACUC #16-175).

**Table 1.** Participant demographics.

| **Table 1.** Participant Demographics. |  |  |
| --- | --- | --- |
|  | 1-2-3 Treat | Treat Hide & Seek |
|  | **N = 5,596<sup>†</sup>** | **N = 3,775<sup>†</sup>** |
| Dog Sex |  |  |
| female | 2,845 (51%) | 1,915 (51%) |
| male | 2,751 (49%) | 1,860 (49%) |
| Dog weight (kg) | 23 (12); 1-96 | 23 (12); 2-88 |
| Dog age (years) | 7.1 (3.7); 0.5-23.0 | 7.3 (3.5); 0.8-23.5 |
| Owner age (years) |  |  |
| 18-24 | 63 (1.1%) | 56 (1.5%) |
| 25-34 | 559 (10.0%) | 364 (9.6%) |
| 35-44 | 689 (12%) | 446 (12%) |
| 45-54 | 942 (17%) | 629 (17%) |
| 55-64 | 1,621 (29%) | 1,113 (29%) |
| 65-74 | 1,483 (27%) | 1,004 (27%) |
| 75 and older | 239 (4.3%) | 163 (4.3%) |
| Owner education |  |  |
| No schooling completed | 1 (<0.1%) | 0 (0%) |
| Nursery school to 8th grade | 0 (0%) | 0 (0%) |
| Some high school, no diploma | 6 (0.1%) | 2 (<0.1%) |
| High school graduate | 94 (1.7%) | 49 (1.3%) |
| High school equivalent degree (such as GED) | 13 (0.2%) | 10 (0.3%) |
| Some college credit, no degree | 376 (6.7%) | 233 (6.2%) |
| Trade, technical, or vocational training | 124 (2.2%) | 78 (2.1%) |
| Associate degree | 280 (5.0%) | 181 (4.8%) |
| Bachelor's degree | 1,887 (34%) | 1,269 (34%) |
| Master's degree | 1,703 (30%) | 1,173 (31%) |
| Professional degree (such as DVM, MD, DDS, or JD) | 548 (9.8%) | 393 (10%) |
| Doctorate degree (such as PhD, DrPH, or DPhil) | 564 (10%) | 387 (10%) |
| Hispanic or Latino | 175 (3.1%) | 108 (2.9%) |
| Owner Race |  |  |
| White | 5,348 (96%) | 3,606 (96%) |
| Black or African American | 46 (0.8%) | 31 (0.8%) |
| Asian | 206 (3.7%) | 151 (4.0%) |
| American Indian | 53 (0.9%) | 32 (0.8%) |
| Alaska Native | 1 (<0.1%) | 0 (0%) |
| Native Hawaiian | 5 (<0.1%) | 2 (<0.1%) |
| Pacific Islander | 15 (0.3%) | 7 (0.2%) |
| Other | 101 (1.8%) | 58 (1.5%) |
| <sup>†</sup> n (%); Mean (SD); Min-Max |  |  |

### Procedure

In both cognitive tasks, community scientists (hereafter, participants) followed tutorials with printable and video-based instructions covering the required materials, task setup, and implementation. Both cognitive tasks required the hiding of food rewards in small containers (∼ 10 x 10 x 6cm) at marked locations in the testing area. Participants received instructions and printable templates for constructing these containers using scrap cardboard, or other common household materials (e.g., tissue boxes or milk cartons). Following set up of the testing area (measuring and marking locations described below), participants followed an interactive tutorial with trial-by-trial instructions and electronic data entry using the Research Electronic Data Capture (REDCap) platform (Harris et al., 2009). If needed, participants had access to printable instructions and a scoring sheet, from which data could be entered later.

### 1-2-3 Treat

#### Overview

In this activity, three boxes were positioned in an array. A handler walked the dog to each of the boxes in sequence. At each location, the handler visibly baited the box, and at two of the three locations the dog was allowed to immediately consume the reward. Following the baiting procedure, the dog was released from the start position, equidistant from the three boxes, to search for the sole remaining reward. Accuracy of the first location searched was recorded across 9 trials.

### Activity setup and warm-up trials

Participants were instructed to mark a start location from which their dog would begin each trial, and three additional locations, each 1.82m from the start location, where the boxes containing food rewards would be positioned (Figure 1). Participants were then guided through three warm-up trials in which dogs gained practice searching for food rewards at each of the three locations used in subsequent test trials. On these trials (one trial per location), the handler placed one box on the target location, walked the dog directly to it, placed a small food reward into the box, walked the dog back to the start position and released the dog to search for the reward. If the dog did not independently find the reward within 20s, the handler led the dog to it. If the dog did not find the food independently in at least 2 of the 3 warm-up trials, handlers were instructed to conduct an additional set of 3 warm-ups before advancing to test trials.

**Figure 1.**
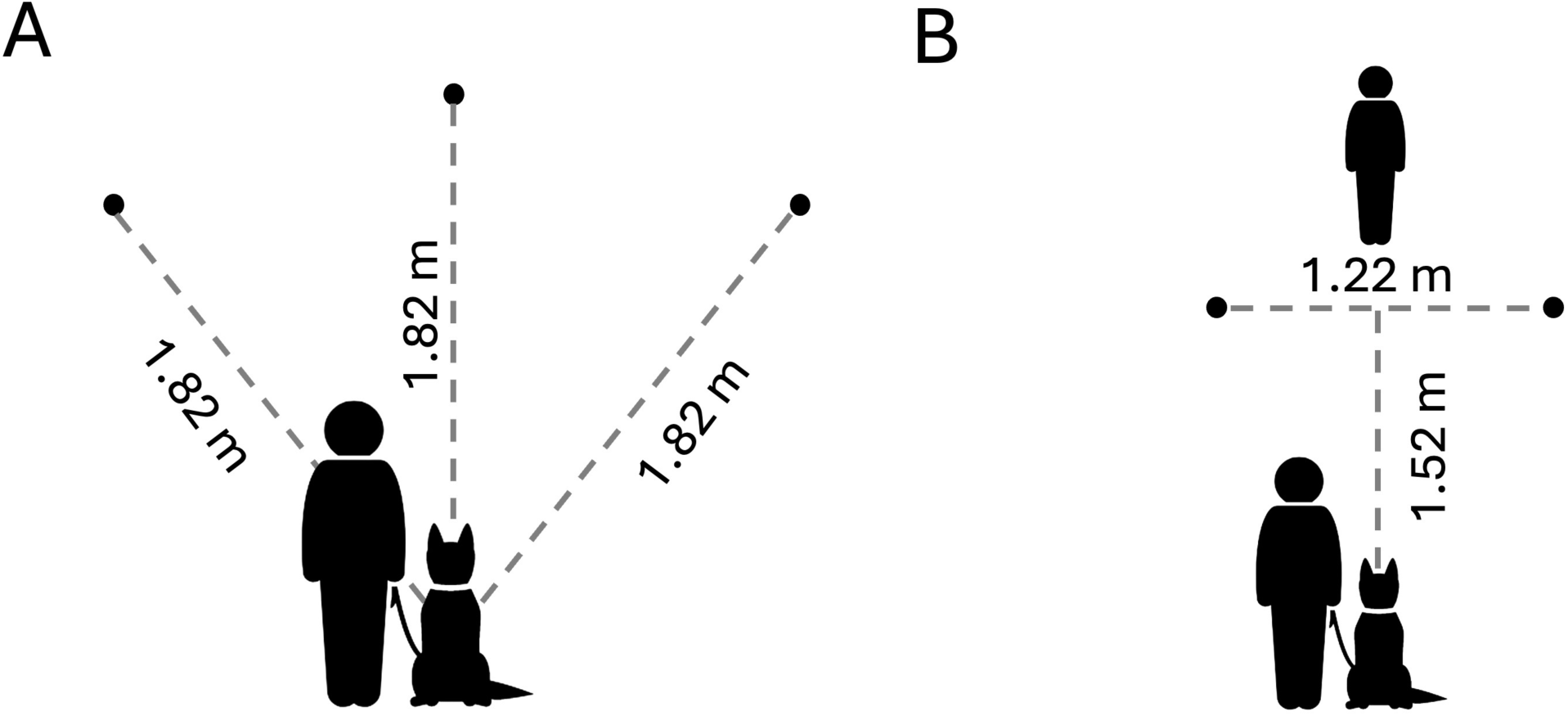
Spatial layout for the (A) 1-2-3 Treat, and (B) Treat Hide & Seek tasks. Circles indicate the locations where food was hidden, relative to the dog’s starting positions. Not to scale.

### Test trials

Prior to test trials, participants selected which side of their body they would handle the dog on and this determined whether they walked the dog in a clockwise (dog on handler’s left) or counterclockwise (dog on handler’s right) pattern. This element of the protocol was implemented to facilitate the handler preventing the dog from immediately consuming the reward during baiting by standing between the dog and the box in which the reward was placed. On each trial, the handler and dog began at the start position, walked a circular pattern around the boxes, visiting each box in sequence, and returned to the start position where the dog was released to search.

At each box, the handler visibly placed a small food reward into the box. At two of the three locations the dog was allowed to immediately consume this reward, whereas at one location the handler prevented the dog from obtaining the reward by immediately walking past the box following its baiting. The location at which the dog was initially prevented from obtaining the reward was balanced across trials (3 trials per location). Upon release from the start line, dogs were allowed 20s to search for the reward, and the first location searched (defined as the dog’s snout entering the area above the box) was recorded. Participants were instructed to let their dog search in only one box per trial. If the dog did not choose correctly, the participant was instructed to prevent further searching using the leash. If appropriate, participants could also select response options indicating that (a) the dog did not search in any box, (b) something went wrong, or (c) the dog was not enjoying the activity and data collection needed to be discontinued.

### Treat Hide & Seek

#### Overview

In this activity, one of two boxes positioned in front of the dog was visibly baited by one person (the experimenter). Following a variable delay (0, 10, 20, or 40s), a second person (the handler) released the dog to search for the reward. Accuracy of the first search was recorded across a series of 16 trials (4 trials per delay duration).

### Activity setup and warm-up trials

Participants were instructed to mark a start location from which their dog would begin each trial (held on leash by a second person acting as handler), two additional locations for the boxes 1.52m in front of and 0.61m to the left and right sides of the start location, and a location 0.3m beyond and centered between the box locations for the experimenter (Figure 1). In warm-up trials the experimenter placed a single box at the left or right location (balanced across warmups) and baited the box with a small food reward as the dog watched from the start position. Immediately following baiting, the experimenter looked down, started a stopwatch and called “okay”, cueing the handler to release the dog to search for the food. The experimenter recorded whether the dog obtained the food independently within 20s on each trial. Two trials were conducted at each location.

### Test trials

In test trials, both boxes were positioned on their respective marks in front of the dog. Dogs watched as the experimenter baited one of the boxes. The experimenter then looked down, initiated the stopwatch and called “okay” for the handler to release the dog once the specified delay had elapsed. Experimenters were instructed to continue gazing down at the stopwatch to avoid cuing the dog inadvertently and to let the dog search for no longer than 20s. At the conclusion of each trial, the experimenter recorded which box the dog searched first (defined as the snout entering the area above the box). Dogs were not prevented from searching in more than one box, and could search until they found the treat, but only the first location searched was scored. If appropriate, participants could also select response options indicating that (a) the dog did not search in any box, (b) something went wrong, or (c) the dog was not enjoying the activity and data collection needed to be discontinued. The delay before dogs were released to search increased across blocks of four trials (trials 1-4 = 0s delay; trials 5-8 = 10s, trials 9-12 = 20s, trials 13-16 = 40s) and the correct location was balanced within block.

### Analysis

#### Dependent measures

For 1-2-3 Treat (hereafter 123T), we used the overall percentage of correct responses as our primary summary score. Only trials on which the dog searched in one of the containers were included in this score. We retained observations for analysis if there were a minimum of 6 trials in which the dog searched in one of the boxes (94% of observations). For secondary models of accuracy as a function of the correct location, we filtered data to cases in which all 3 trials at a given location had a valid search (85% of observations). Treat Hide & Seek (hereafter THS) summary scores were similarly calculated as the overall percentage of correct responses across trials in which dogs searched in one of the containers. We retained THS summary scores for analysis if a dog made valid choices on 12 or more of the 16 trials (96% of observations). For secondary models of accuracy as a function of delay length, we filtered data to observations with complete data for all delay durations (95% of observations).

### Statistical Models

To estimate associations between age and cognitive scores we fit linear models incorporating a second-order orthogonal polynomial term for age, with covariates for sex, and body weight (standardized to have a mean of 0 and standard deviation of 1). To assess effects of the spatial location of the remaining reward (123T) or delay length (THS) we used similar linear mixed models with an additional term for reward location or delay length, and a random intercept for the dog identifier. Post-hoc comparisons from these models incorporated Tukey’s method for adjustment of p values. To facilitate interpretation of the non-linear effect for age, we report marginal estimates of performance for dogs of different ages (Arel-Bundock et al., 2024). To test the hypothesis that associations between age and cognitive performance are moderated by body weight (a proxy for expected lifespan), we compared the base models described above to alternative models incorporating an interaction between the polynomial term for age and dog body weight (owner report). We assessed the relative support for each model using Bayes factors based on BIC approximation, with interpretation guided by Jeffreys’ scale (Jeffreys, 1998; Kass & Raftery, 1995). As secondary analyses, we fit similar models using a genomic prediction of dog height as our measure of size (supplemental material; Dog Aging Project, 2025)

To compare data gathered by community scientists in the home environment with data collected by trained experimenters in professional research settings, we used a matching strategy. For each of the dogs tested by research professionals, 5 matches from the community science dataset were included based on nearest neighbor propensity score matching using logistic regression, as implemented in the MatchIt R package (Ho et al., 2011). Matching criteria included age (at time of test), body mass, and sex. We included calipers (maximum acceptable distance for the match) of 1 year for age and 2 kg for body mass. Following matching, standardized groups mean differences for age and weight were < 0.1. Using these data, we fit linear models with the overall task score modeled as a function of a second order polynomial term for age, sex, weight, and the data collection context (community science vs professional administration). We used the marginaleffects R package (Arel-Bundock et al., 2024) for g-computation in the matched sample to estimate the average treatment effect in the treated (ATT). These estimates used cluster-robust standard errors with matching stratum membership as the clustering variable.

## Results

### 1-2-3 Treat

On average, dogs first searched in the correct location on 68.8 ± 0.3% of trials, significantly higher than chance expectation (one-sample t-test, t_5237_ = 116.95, p < 0.01). Overall performance varied significantly as a function of age (Figure 2; Table 2). On average, a 3-year-old dog responded correctly on 72% of trials whereas a 13-year-old dog was correct on 62% of trials. Accuracy also varied in relation to the remaining reward’s order in the baiting sequence (𝜒^2^_2_ = 1209.6, p < 0.01). Performance was best when the reward was at the final baiting location (82.7 ± 0.5%), intermediate when it was at the first baiting location (66.5 ± 0.5%), and worst at the middle baiting location (62 ± 0.5%), consistent with a serial position effect (Bolhuis & Van Kampen, 1988; Smyth & Scholey, 1996). Poorer performance at older ages was observed at all locations with the effect of age estimated to be more linear when the reward was at the first and second locations in the baiting sequence (Table 2, Figure 2). A model incorporating an interaction between age and body weight estimated highly similar associations with age in smaller and larger dogs (Figure 2) and a comparison of this model to one without the interaction yielded decisive evidence in favor of the latter (Bayes factor = 2,890).

**Figure 2.**
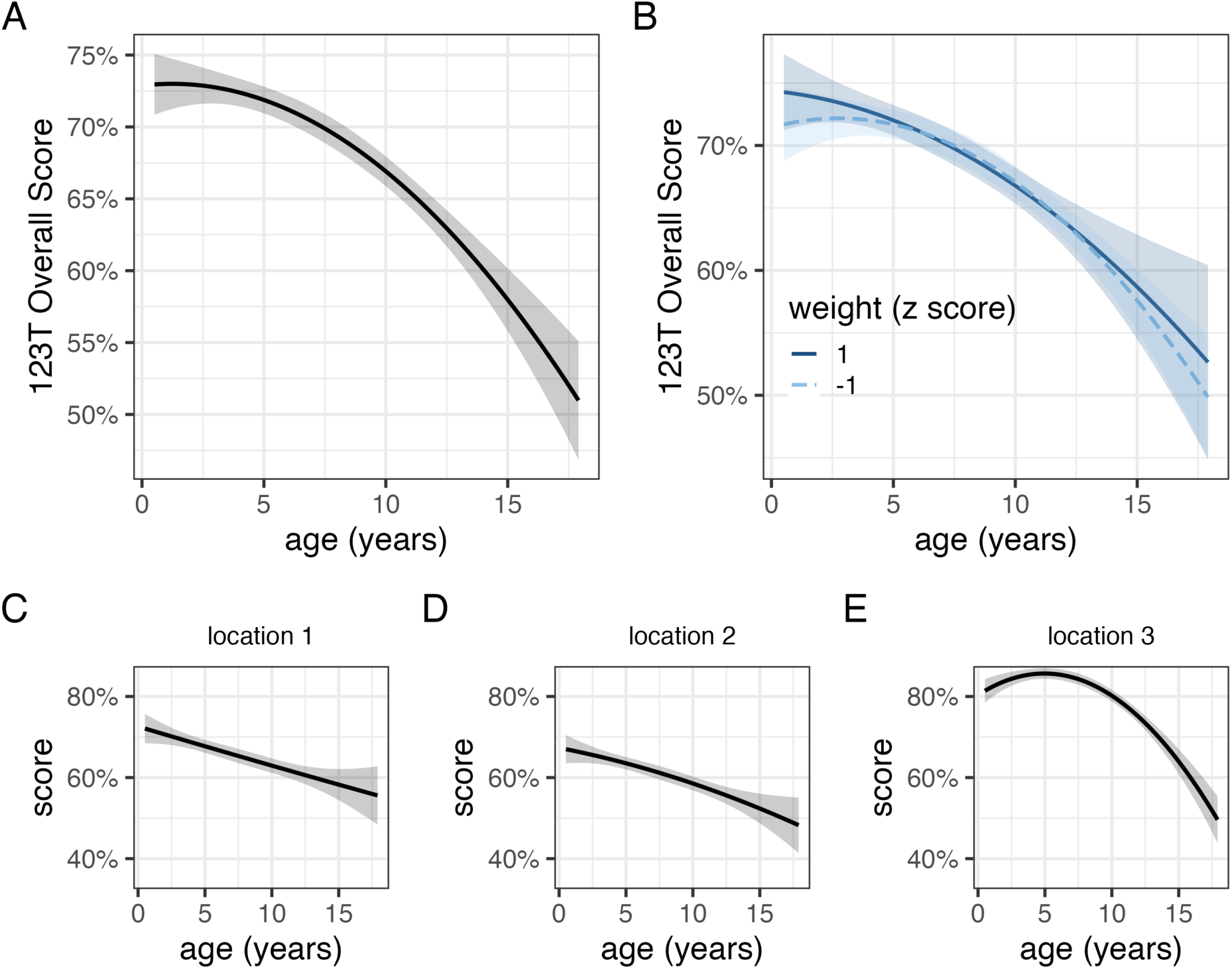
Relationships between age and performance on 1-2-3 Treat. The relationship with age (A) was highly similar across dogs of different body masses (B). Impaired performance at older ages was observed at all locations in the baiting sequence (C,D,E).

**Table 2.** Statistical models for 1-2-3 Treat.

| 1-2-3 Treat Models |  |  |  |  |  |  |  |  |  |  |  |  |
| --- | --- | --- | --- | --- | --- | --- | --- | --- | --- | --- | --- | --- |
|  | overall score |  |  | location 1 |  |  | location 2 |  |  | location 3 |  |  |
|  | Beta | 95% CI | p-value | Beta | 95% CI | p-value | Beta | 95% CI | p-value | Beta | 95% CI | p-value |
| age |  |  |  |  |  |  |  |  |  |  |  |  |
| age | -277 | -320, -234 | <0.001 | -227 | -295, -160 | <0.001 | -251 | -318, -184 | <0.001 | -319 | -378, -261 | <0.001 |
| age<sup>2</sup> | -82 | -125, -39 | <0.001 | 1.8 | -66, 69 | >0.9 | -23 | -90, 44 | 0.5 | -219 | -277, -161 | <0.001 |
| sex |  |  |  |  |  |  |  |  |  |  |  |  |
| female | — | — |  | — | — |  | — | — |  | — | — |  |
| male | -0.98 | -2.2, 0.21 | 0.11 | -1.1 | -3.2, 0.91 | 0.3 | -1.1 | -3.0, 0.91 | 0.3 | -0.76 | -2.4, 0.88 | 0.4 |
| weight (z score) | 0.18 | -0.44, 0.79 | 0.6 | 0.32 | -0.75, 1.4 | 0.6 | -0.43 | -1.4, 0.59 | 0.4 | 0.09 | -0.77, 0.94 | 0.8 |
| Abbreviation: CI = Confidence Interval |  |  |  |  |  |  |  |  |  |  |  |  |

### Treat Hide & Seek

On average, dogs searched in the correct location on 92% of trials, significantly higher than chance expectation (one-sample t-test, t_3629_ = 205.1, p < 0.01). THS performance varied significantly by age (Figure 3; Table 3). On average, a 3-year-old dog scored ∼93% whereas a 13-year-old dog scored ∼88%. Accuracy varied in relation to the delay length before dogs were allowed to search, but surprisingly performance was modestly better at longer delays (β_minute_ = 0.8% ± 0.4%, 𝜒^2^_1_ = 4.03, p = 0.04). Poorer performance for older dogs was observed at all delay lengths with older dogs estimated to perform worst at the two longest delays (20s & 40s). A model with an interaction between age and body weight estimated similar associations between performance and age in small and large dogs (Figure 3), and again, model comparison strongly favored the model without the weight × age interaction (Bayes factor = 243).

**Figure 3.**
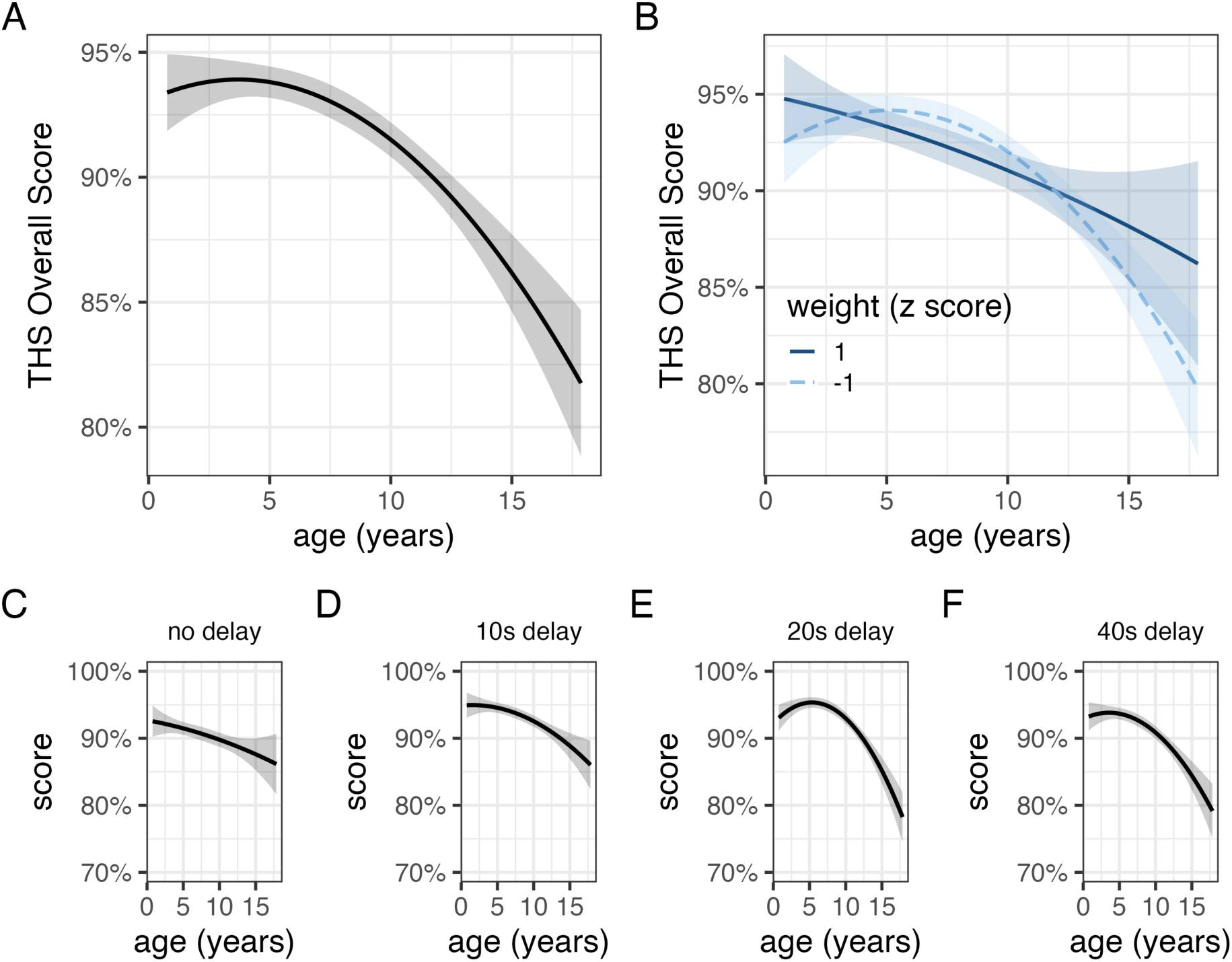
Relationships between age and performance on Treat Hide & Seek. The relationship with age (A) was similar across dogs of different body masses (B). Impaired performance at older ages was observed across delay intervals (C,D,E,F).

**Table 3.** Statistical models for Treat Hide & Seek.

| Treat Hide & Seek Models |  |  |  |  |  |  |  |  |  |  |  |  |  |  |  |
| --- | --- | --- | --- | --- | --- | --- | --- | --- | --- | --- | --- | --- | --- | --- | --- |
|  | overall score |  |  | no delay |  |  | 10s delay |  |  | 20s delay |  |  | 40s delay |  |  |
|  | Beta | 95% CI | p-value | Beta | 95% CI | p-value | Beta | 95% CI | p-value | Beta | 95% CI | p-value | Beta | 95% CI | p-value |
| age |  |  |  |  |  |  |  |  |  |  |  |  |  |  |  |
| age | -108 | -132, -84 | <0.001 | -72 | -108, -35 | <0.001 | -88 | -117, -59 | <0.001 | -118 | -147, -88 | <0.001 | -127 | -159, -95 | <0.001 |
| age<sup>2</sup> | -48 | -72, -24 | <0.001 | -7.4 | -44, 29 | 0.7 | -26 | -55, 3.1 | 0.080 | -85 | -114, -55 | <0.001 | -55 | -87, -23 | <0.001 |
| sex |  |  |  |  |  |  |  |  |  |  |  |  |  |  |  |
| female | — | — |  | — | — |  | — | — |  | — | — |  | — | — |  |
| male | -0.96 | -1.8, -0.16 | 0.019 | -0.89 | -2.1, 0.33 | 0.2 | -0.57 | -1.6, 0.41 | 0.3 | -1.1 | -2.1, -0.08 | 0.035 | -0.90 | -2.0, 0.20 | 0.11 |
| weight (z score) | -0.29 | -0.70, 0.13 | 0.2 | -0.56 | -1.2, 0.07 | 0.081 | -0.14 | -0.65, 0.37 | 0.6 | 0.01 | -0.51, 0.53 | >0.9 | -0.45 | -1.0, 0.12 | 0.12 |
| Abbreviation: CI = Confidence Interval |  |  |  |  |  |  |  |  |  |  |  |  |  |  |  |

### Matching analysis

For 123T, scores obtained by community scientists were on average 4.6% higher than those for dogs tested using conventional methods, a difference that was not statistically significant (z = - 0.98, p = 0.33). Both datasets were characterized by a similar pattern in which performance was highest when the sole remaining treat was in the last location baited, compared to when it was in the 1^st^ or 2^nd^ location baited (Figure 4). For THS, we observed similar scores in matched dogs tested by community scientists and trained experimenters. On average, scores were 2.6% higher in the group tested by trained experimenters, a difference that was not statistically significant (z = 1.18, p = 0.24). In the conventional dataset, average scores decreased monotonically with delay length, but this pattern was not observed in the matched community science sample (Figure 4).

**Figure 4.**
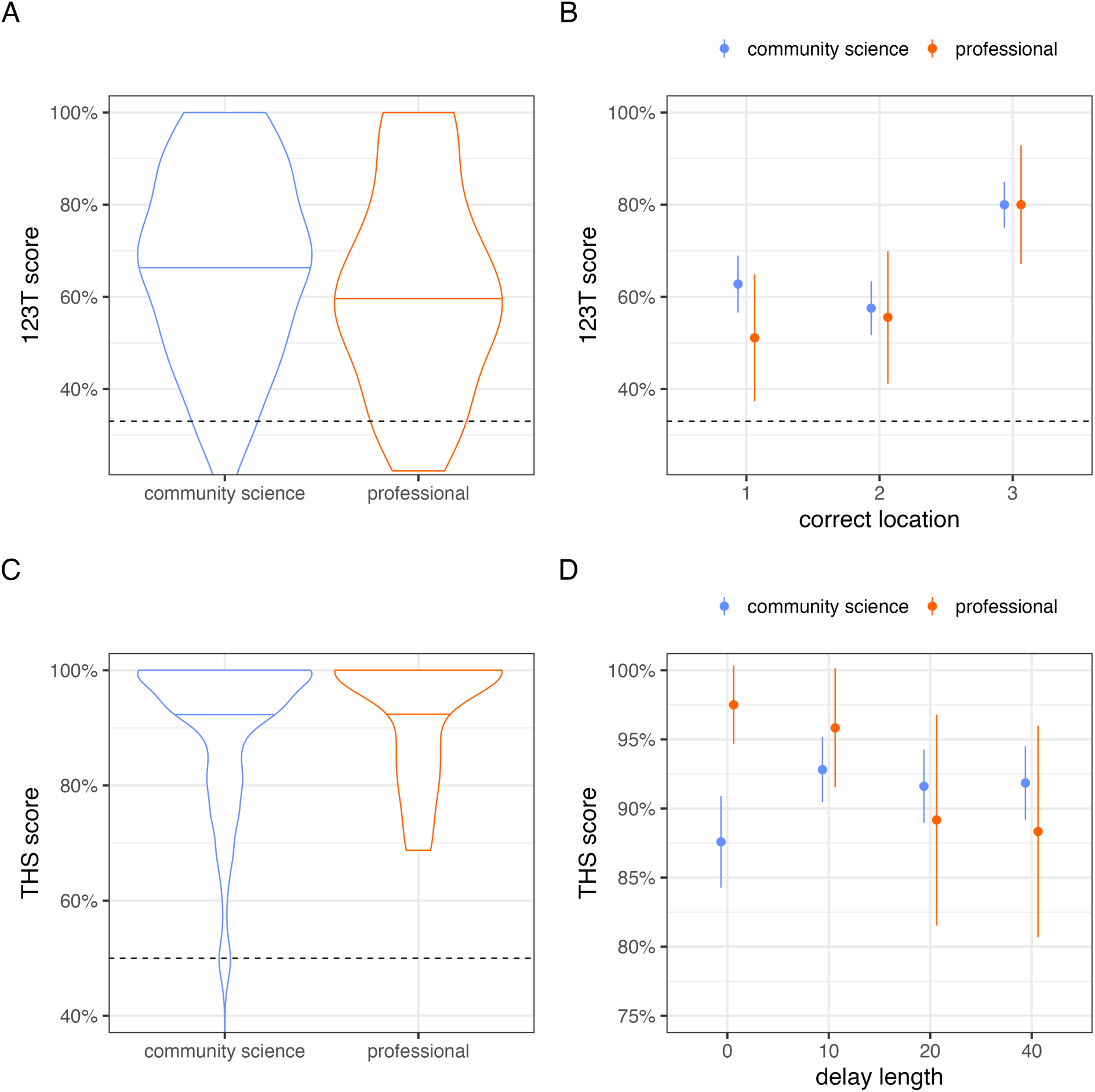
Comparison of data collected by community scientists and research professionals. (A) On 1-2-3 Treat, overall scores were similar between datasets, and (B) both datasets were characterized by a serial position effect. (C) On Treat Hide & Seek, scores were similar, and near ceiling, in both datasets. (D) A trend toward worse performance at longer delays was observed in data collected by research professionals but not community scientists. Points and line intervals in panels B and D represent means and 95% confidence intervals.

## Discussion

Large scale studies of dog cognition face the challenges of administering objective cognitive assessments to nonverbal organisms. We designed two novel cognitive assessments of spatial memory, which were administered by community scientists in a sample of thousands of companion dogs throughout the United States, as a part of the Dog Aging Project. Both tasks demonstrated age-related variation in performance, and sensitivity to age-related cognitive changes was a central goal in developing these measures. Community scientists generated data that were similar to those obtained by research professionals, consistent with previous studies in which dog owners administered simple cognitive tests at home (Hoel et al., 2021; Stewart et al., 2015). Using these data we tested the hypothesis that relationships between age and cognitive function are moderated by body weight, as a proxy for expected lifespan in dogs. We found strong evidence against this hypothesis, instead observing highly similar relationships between age and performance across dogs of different sizes. This finding is consistent with previous cross-sectional analyses that have estimated this relationship using data from laboratory-administered tests (Hargrave et al., 2024), community science-based assays (Watowich et al., 2020), and clinically-validated dementia scales (Azkona et al., 2009; Salvin et al., 2010; but see Turcsán & Kubinyi, 2024). Given that large dogs are shorter-lived and exhibit accelerated aging across a diverse spectrum of characteristics, these findings raise the possibility that large size might confer a protective effect for brain and cognitive aging.

We hypothesize that large body size may be associated with healthy brain and cognitive aging for the following two reasons. First, variation in dog size is driven predominantly by insulin-like growth factor 1 (IGF-1) signaling (Hoopes et al., 2012; Sutter et al., 2007). In the brain, IGF- 1 promotes neurogenesis and synaptogenesis, protects neurons from chemical toxicity, and contributes to β-amyloid clearance, which are all mechanisms that may prolong cognitive health (Carro et al., 2002; Sonntag et al., 2005). Although IGF-1 has also been associated with deleterious effects in the central nervous system, and may play opposing roles in brain aging (Gubbi et al., 2018), neuroprotective roles for IGF-1 have been widely recognized and may confer advantages to larger dogs. Second, larger dogs have larger brains that contain a greater number of neurons (Hecht et al., 2019; Jardim-Messeder et al., 2017) which may contribute to cognitive or brain reserve, or the ability to maintain cognitive function despite neurodegeneration (Katzman et al., 1989; Stern, 2002). In human studies neuronal density in the locus coeruleus has been associated with both higher baseline cognitive function and slower cognitive decline (Wilson et al., 2013). Although previous studies have identified positive associations between brain volume and cognitive performance in dogs (Horschler et al., 2019), we did not observe this effect in the current work. Thus, if large brain size alone confers a protective effect, it may result from better maintenance of function, rather than an early life performance advantage.

Our ability to test these, and other hypotheses regarding predictors of variation in dog cognitive decline will depend critically on the ability to accurately assess cognitive function in large and diverse companion dog samples. The community science measures introduced here show promise for this approach with several notable strengths, but also some important limitations. Both 123T and THS were sensitive to age-related variation in performance, a central design goal for these assessments. The overall association with age was similar to that observed in laboratory studies, with peak performance during the first 5 years of life and markedly poorer performance at older ages. Sensitivity to mild, mid-life dysfunction is an important attribute of these measures and contrasts with survey-based assessments of cognitive dysfunction which predominantly capture diminished performance significantly later in life (Yarborough et al., 2022). Another strength of the approach presented here is that data are derived from simple experiments in which dogs are presented with a standardized problem and responses are scored using objectively defined behavioral criteria. This experimental approach avoids common pitfalls associated with survey research, though it is associated with its own limitations, which we discuss below. Lastly, data collected by community scientists were similar to those obtained by research professionals, consistent with previous studies employing community scientists in experimental designs (Stewart et al., 2015).

Despite these strengths, the current approach also has several notable limitations. First, it is unrealistic to expect that members of the public, volunteering their time in a home environment and implementing these procedures once annually will generate data of similar quality to those obtained by trained research professionals working in a laboratory setting. We attempted to accommodate potential errors and unexpected occurrences by providing response options indicating that something went wrong, and enabling participants to discontinue data collection at any point. However, we were unable to directly monitor the fidelity with which protocols were implemented or the accuracy of data entered by participants. Second, although we obtained a large sample relative to those associated with laboratory studies, participation rates in these activities were lower than those for non-experimental, survey-based activities in the Dog Aging Project. The lower participation rate likely reflects the higher demands of gathering this type of data, which required participants to have access to a suitable testing space, a second person to help with data collection, and sufficient time to set up and implement each assessment. We expect that improved methods for engaging and retaining community scientists will be critical for increasing participation rates, and that systematic analysis of factors influencing participation will facilitate advances in this area (e.g. Asingizwe et al., 2020; McAteer et al., 2021; Shinbrot et al., 2023). Lastly, scores on THS exhibited a ceiling effect, in which most dogs made no errors, potentially masking variation among the higher-performing fraction of our sample. This limitation could potentially be remedied by modifications to increase task difficulty, for example through the addition of longer delays, additional search locations, or distraction procedures during the delay (Salomons et al., 2024; Bray et al., 2021).

Collectively, our findings demonstrate the promise of experimental assessments of dog cognition using a community science model. Our analyses of cross-sectional data using this approach demonstrate sensitivity to age-related variation in short-term spatial memory and corroborate findings from previous work suggesting that relationships between age and cognitive function are not moderated by body mass, despite a strong negative correlation between dog size and longevity. Ongoing longitudinal assessment using these measures will support future analyses of within-individual change, enabling powerful tests of whether rates of cognitive decline differ between small and large dogs. Lastly, integration of these data with molecular measures collected in the Dog Aging Project (Creevy et al., 2022; Prescott et al., 2025) will provide a powerful opportunity to test hypotheses about the mechanisms underlying individual differences in cognitive aging.

## Funding and Acknowledgements

This research is based on publicly available data collected by the Dog Aging Project, under U19 grant AG057377 (PI: Daniel Promislow) and R24AG073137 (MPI: Kaeberlein, Keene, McGrath) from the National Institute on Aging, a part of the National Institutes of Health, and by additional grants and private donations, including generous support from the Dogtopia Foundation, the Glenn Foundation for Medical Research, the Tiny Foundation Fund at Myriad Canada, and the WoodNext Foundation. These data are housed on the Terra platform at the Broad Institute of MIT and Harvard. The content is solely the responsibility of the authors and does not necessarily represent the official views of the National Institutes of Health. The authors thank all Dog Aging Project participants for their important contributions to this study.

## Dog Aging Project Consortium authors

Joshua M. Akey^14^, Rozalyn M. Anderson^15,16^, Elhanan Borenstein^17,18,19^, Marta G. Castelhano^20,21^, Amanda E. Coleman^22^, Kate E. Creevy^23^, Matthew D. Dunbar^3^, Virginia R. Fajt^24^, Jessica M. Hoffman^25^, Erica C Jonlin^26,27^, Matt Kaeberlein^26,28^, Elinor K. Karlsson^29,30,^ Kathleen F. Kerr^31^, Jing Ma^32^, Stephanie McGrath^33,34^, Natasha J Olby^35^, May J Reed^36^, Audrey Ruple^37^, Stephen M. Schwartz^38,39^, Sandi Shrager^40^, Noah Snyder-Mackler^41-43^, M. Katherine Tolbert^23^

^14^Lewis-Sigler Institute for Integrative Genomics, Princeton University, Princeton, NJ, USA

^15^University of Wisconsin Madison, Madison, WI

^16^GRECC William S Middleton Memorial Veterans Hospital, Madison WI

^17^Department of Clinical Microbiology and Immunology, Sackler Faculty of Medicine, Tel Aviv University, Tel Aviv, Israel

^18^Blavatnik School of Computer Science, Tel Aviv University, Tel Aviv, Israel

^19^Santa Fe Institute, Santa Fe, NM, USA

^20^Cornell Veterinary Biobank, College of Veterinary Medicine, Cornell University, Ithaca, NY, USA

^21^Department of Clinical Sciences, College of Veterinary Medicine, Cornell University, Ithaca, NY, USA

^22^Department of Small Animal Medicine and Surgery, College of Veterinary Medicine, University of Georgia, Athens, GA, USA

^23^Department of Small Animal Clinical Sciences, Texas A&M University School of Veterinary Medicine & Biomedical Sciences, College Station, TX, USA

^24^Department of Veterinary Physiology and Pharmacology, Texas A&M University School of Veterinary Medicine & Biomedical Sciences, College Station, TX, USA

^25^Department of Biological Sciences, Augusta University, Augusta, GA, USA

^32^Department of Laboratory Medicine and Pathology, University of Washington School of Medicine, Seattle, WA, USA

^27^Institute for Stem Cell and Regenerative Medicine, University of Washington, Seattle, WA, USA

^28^Department of Oral Health Sciences, University of Washington, Seattle, WA, USA

^29^Bioinformatics and Integrative Biology, University of Massachusetts Chan Medical School, Worcester, MA, USA

^30^Broad Institute of MIT and Harvard, Cambridge, MA, USA

^31^Department of Biostatistics, University of Washington, Seattle, WA, USA

^32^Division of Public Health Sciences, Fred Hutchinson Cancer Research Center, Seattle, WA, USA

^33^ Department of Clinical Sciences, College of Veterinary Medicine and Biomedical Sciences, Colorado State University, Ft. Collins, CO

^34^Brain Research Center at Colorado State University

^35^Department of Clinical Sciences, College of Veterinary Medicine, North Carolina State University, Raleigh, NC, USA

^36^Department of Medicine, Division of Gerontology and Geriatric Medicine, University of Washington School of Medicine, Seattle, WA

^37^Department of Population Health Sciences, Virginia-Maryland College of Veterinary Medicine, Virginia Tech, Blacksburg, VA, USA

^38^Epidemiology Program, Fred Hutchinson Cancer Research Center, Seattle, WA, USA

^39^Department of Epidemiology, University of Washington, Seattle, WA, USA

^40^Collaborative Health Studies Coordinating Center, Department of Biostatistics, University of Washington, Seattle, WA, USA

^41^School of Life Sciences, Arizona State University, Tempe, AZ, USA

^42^Center for Evolution and Medicine, Arizona State University, Tempe, AZ, USA

^43^School for Human Evolution and Social Change, Arizona State University, Tempe, AZ, USA

## Author Contributions

Conceptualization: SHH, ELM, DELP, DAP Consortium

Methodology: SHH, AJK, TAB, EEB, LELCD, BSK, ELM

Software: SHH, ELM

Formal Analysis: SHH, ELM

Investigation: SHH, EK, ELM, GEA, DAR, EEB, AJK, DELP, DAP Consortium

Resources: DAP Consortium

Data Curation: DAP Consortium

Writing – Original Draft: SHH, AJK, ELM

Writing – Review & Editing: All authors

Visualization: SHH, ELM

Supervision: ELM

Project Administration: AJK, DAP Consortium

Funding Acquisition: DELP, ELM, GEA, DAR, DAP Consortium

