## supplemental material for "Functional assessments of short-term spatial memory in the Dog Aging Project identify strong associations with age that are not moderated by body mass"

The size distribution of dogs that participated in the cognitive tasks was similar to that for non-participating dogs (Figure S1A). Participant dogs were moderately younger than non-participant dogs (mean ± SD age, participants = 5.2 ± 3.5 years, non-participants = 7.0 ± 4.4 years; Figure S1B). Lastly, owners of participating dogs tended to be moderately older than those of non-participant dogs (Figure S1C).

**Figure S1.** Comparison of demographic characteristics among dogs/owners that participated in the cognitive tasks, to those that did not.  Histograms in panels A and C use bins following data conventions in the Dog Aging Project. Bar heights reflect density to facilitate comparison of participant and non-participant distributions. Data in panel B reflect the dog's age at the time of enrollment. Data in panel C reflect the owner's age category at time of enrollment. Non-participant data for all comparisons was restricted to dogs that enrolled during the enrollment date range of participant dogs.

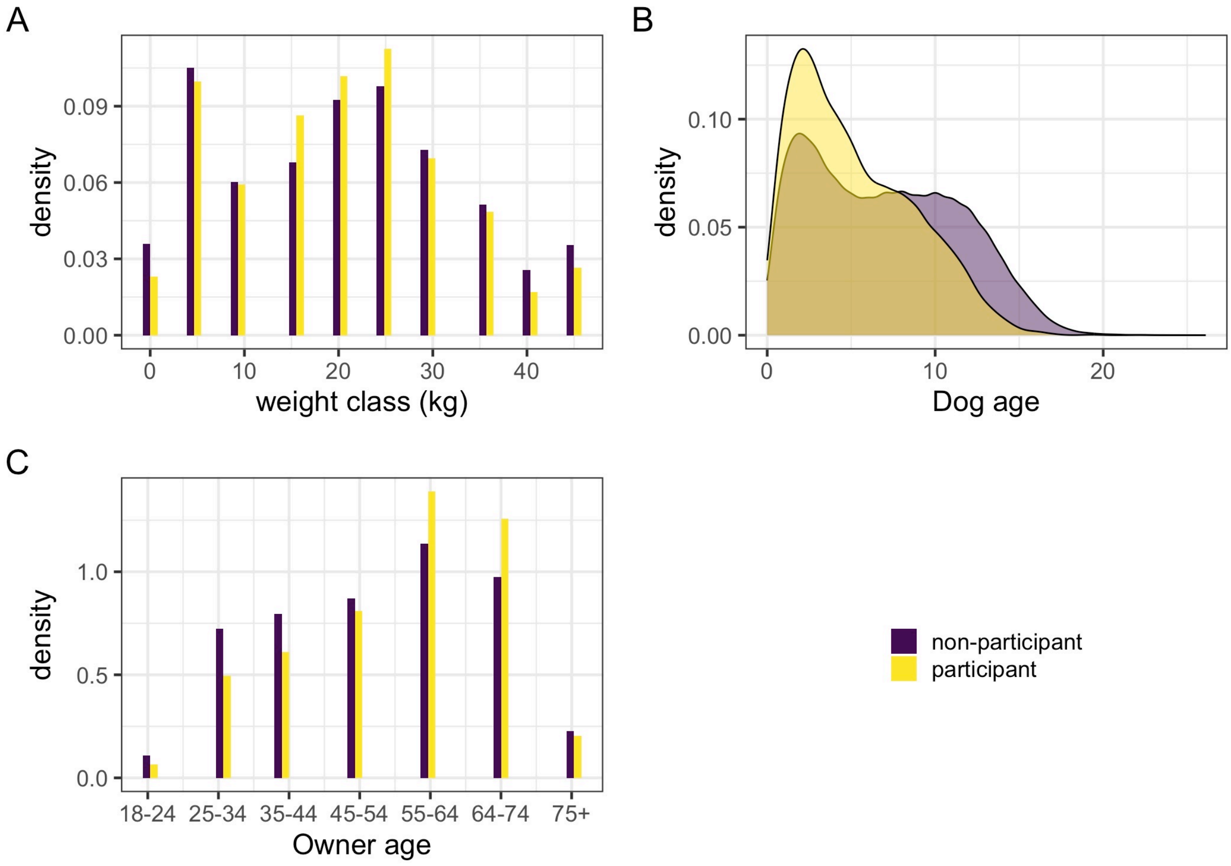

Secondary models using genomic predictions for dog height rather than body weight as a measure of size yielded similar results to our primary models incorporating body weight (Figure S2). For both cognitive tasks model comparison provided substantially strong support for a model without a predicted size × age interaction (123T, bayes factor = 1,870; THS, bayes factor = 867.0). Genomic size predictions for these models were based on a random forest model developed on the Darwin's Ark (darwinsark.org) data set of 1,730 dogs, as described in the Dog Aging Project 2024 data release (Dog Aging Project, 2025).

**Figure S2.** Estimated effects of age at genomically predicted heights 1 SD above and below the sample mean.

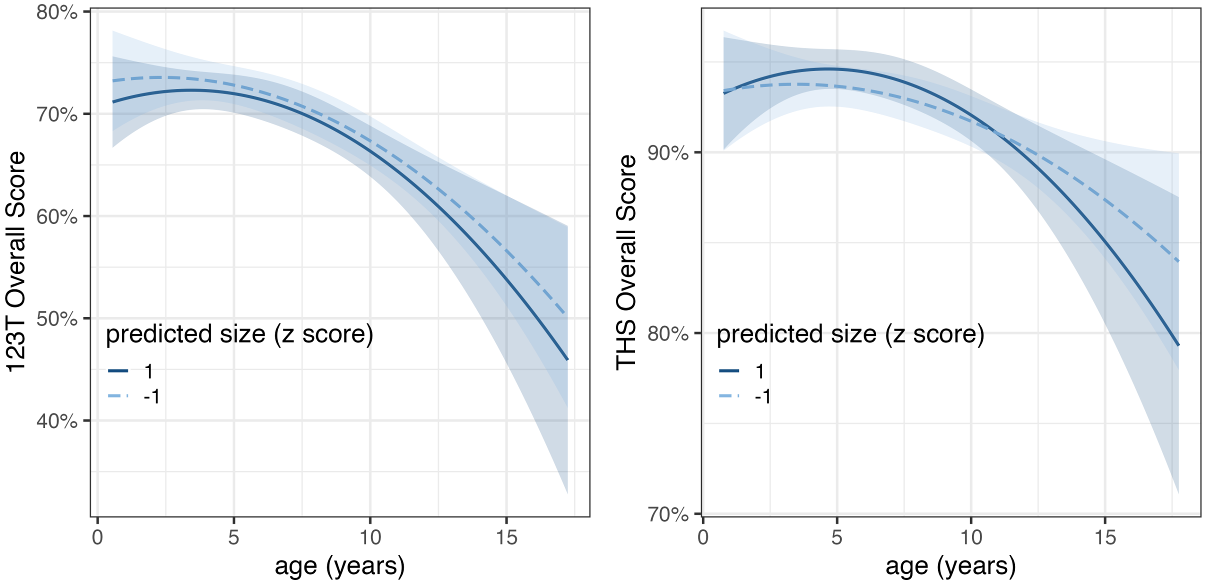
